# Chromosome-encoded IpaH ubiquitin ligases indicate non-human pathogenic *Escherichia*

**DOI:** 10.1101/2021.10.19.464960

**Authors:** Natalia O. Dranenko, Maria Tutukina, Mikhail S. Gelfand, Fyodor A. Kondrashov, Olga O. Bochkareva

## Abstract

Until recently, *Shigella* and enteroinvasive *Escherichia coli* were thought to be primate-restricted pathogens. The base of their pathogenicity is the type 3 secretion system (T3SS) encoded by the *pINV* virulence plasmid, which facilitates host cell invasion and subsequent proliferation. A large family of T3SS effectors, E3 ubiquitin-ligases encoded by the *ipaH* genes, have a key role in the *Shigella* pathogenicity through the modulation of cellular ubiquitination that degrades host proteins. However, recent genomic studies identified *ipaH* genes in the genomes of *Escherichia marmotae,* a potential marmot pathogen, and an *E. coli* extracted from fecal samples of bovine calves, suggesting that non-human hosts may also be infected by these strains, potentially pathogenic to humans.

We performed a comparative genomic study of the functional repertoires in the *ipaH* gene family in *Shigella* and enteroinvasive *Escherichia* from human and predicted non-human hosts. We found that fewer than half of *Shigella* genomes had a complete set of *ipaH* genes, with frequent gene losses and duplications that were not consistent with the species tree and nomenclature. Non-human host IpaH proteins had a diverse set of substrate-binding domains and, in contrast to the *Shigella* proteins, two different types of the NEL C-terminal domain. Only the *ipaH9.8* gene was found in *Escherichia* derived from both human and non-human hosts. These results provide a framework for understanding of *ipaH*-mediated host-pathogens interactions and suggest a need for a genomic study of fecal samples from diseased animals.

## Introduction

Shigellosis is a widespread human intestinal infection disease. Its causative agent, *Shigella,* is one of *Escherichia coli* pathovars, but the genus name is maintained due to medical importance (1,2). Based on the symptoms and molecular features of the infection, the *Shigella* genus has been classified into four species (3). However, these *Shigella* species are not monophyletic and have arisen independently from different non-pathogenic *E. coli* by acquiring a large plasmid that encodes a substantial number of virulence genes (1). *Shigella* and enteroinvasive *E. coli* (EIEC) enter epithelial cells of the colon, multiply within them, and move between adjacent cells (4). Both *Shigella* and EIEC become invasive by acquiering a *pINV* plasmid with essential virulence determinants, including genes encoding the type III secretion system (T3SS) (4). Furthermore, the pathogens’ genomes feature other genomic markers of adaptation to the intracellular lifestyle, such as chromosomal pathogenicity islands, accumulation of mobile elements, and lack of genes coding for bacterial motility or lactose fermentation (5). Because EIEC retain the ability to live outside the host cells and their genomes harbor significantly less mobile elements and pseudogenes, EIEC are believed to be the precursors for *Shigella* lineages (6).

Encoded in the “entry region” of *pINV,* the T3SS proteins have a range of diverse functions, being structural proteins, chaperones that protect *Shigella* and EIEC virulence proteins from aggregation and degradation, and effector proteins that are secreted into the host cell and selectively bind particular host proteins to regulate the host biological activity (7). Numerous pathogenic bacteria affect the ubiquitination pathway of the host. In *Shigella*, novel E3 ubiquitin-ligases encoded by the *ipaH* genes modulate cellular ubiquitination leading to the degradation of host proteins (8). The IpaH proteins are comprised of two domains, the highly conserved, **n**ovel **E**3 **l**igase (NEL) C-terminal domain that binds ubiquitin, and the variable **l**eucine-**r**ich **r**epeat-containing (LRR) N-terminal domain that binds various human proteins hence providing the substrate specificity (9). The IpaH proteins are thought to trigger cell death and to modulate host inflammatory-related signals during bacterial infection; however, the substrate specificity of many IpaH proteins remains uncertain (10,11).

Expression of the *ipaH* genes can be regulated by several transcription factors. MxiE, a transcription activator encoded in the “entry” region of *pINV*, regulates the intracellular expression of genes encoding numerous factors secreted by the type III secretion system, including OspB, OspC1, OspE2, OspF, VirA, and IpaH (12). Two plasmid-encoded virulence transcription factors, VirF and VirB, are known to turn on the *Shigella* virulence by activating major determinants, and thus may also control the *ipaH* genes (13). Both the *virF* and *virB* genes have sites for thermal sensing, and at 30°C both of them are negatively controlled by the global transcriptional silencer H-NS (13,14). Upon invasion of the host organism, H-NS detaches from DNA, switching on the virulence cascades. H-NS normally binds A/T-rich elements making bridges or loops that affect transcription from the target promoters (15,16). The regulatory regions of many virulence genes in *Shigella* have A+T rich tracks and H-NS-bound A+T tracks are common features of mobile elements or prophages, and may be a footprint of a recent horizontal gene transfer (17).

Although naturally *Shigella* was thought to be a primate-specific pathogen, experiments showed that it can infect other animals, yet with lower efficacy (18,19). Recently, *Shigella*-like T3SS and associated effectors were found in *Escherichia marmotae,* a potential invasive pathogen of marmots (20), which was also shown to be able to invade human cells (20). *Shigella* marker genes were also found in isolates obtained from the excrement of bovine calves with diarrhea, although genome-wide data was lacking (21).

Here, we applied a computational approach to predict whether some *Escherichia* may also be an infectious agent of non-human hosts, which, therefore, may serve as a reservoir of human pathogens and virulence genes. For that, we performed a comparative genomic analysis of the *ipaH* genes in *Shigella*, EIEC strains, and putatively invasive *Escherichia* species extracted from non-human hosts. We classified and compared members of the *ipaH* gene family based on domain sequence similarity, genomic location, and positioning of regulatory elements in upstream gene regions. Furthermore, for *Shigella* lineages we reconstructed the evolution of the *ipaH* genes on the species phylogenetic tree revealing multiple gene losses, paralogizations, and horizontal gene transfer.

## Methods

### Dataset of genomes

We downloaded 130 complete genomes of *Shigella* available in GenBank (22) as of November 2020 and three complete genomes of enteroinvasive *Escherichia coli* **(*Supplementary Table S1*)**. Additionally we downloaded all *Escherichia* assemblies extracted from non-human hosts that contained BLAST (23) hits of the NEL-domain of *Shigella* IpaH ***(Supplementary Table S2)***.

### Identification of the ipaH genes

Using pBLAST search of the NEL-domain (PDB: *Shigella flexneri* Effector IpaH1880 5KH1 https://www.rcsb.org/structure/5KH1), we found 445 protein sequences belonging to the E3 ubiquitin-ligase family. Then we clustered the sequences using CD-hit (24) with a threshold of 90% aa identity and performed additional tBLASTn search of representative sequences from each cluster. It allowed us to add 419 sequences including non-annotated genes and pseudogenes. In total, we found and classified 864 *ipaH* sequences **(*Supplementary Table S3*)**. The *ipaH* genes in non-human *Escherichia* were found using the same pipeline and collected in ***Supplementary Table S4***.

### Heatmaps

Heatmaps for sequence similarity were drawn using R packages `seqinr`, `RColorBrewer`, and `gplots`.

### Phylogenetic tree

For construction of the *Shigella* species tree, we used the `PanACoTA` tool (25). It annotates coding regions, finds orthologous groups, and constructs the phylogenetic tree for a concatenated alignment of single-copy common genes. The orthologous groups were constructed with a threshold of 80% aa identity, the phylogenetic tree was constructed with the `IQ-TREE` 2 module (26). The tree was visualised using online `iTOL` (27).

### Annotation of regulatory elements in upstreams

Alignments of the *ipaH* regulatory regions were constructed with the `Pro-Coffee` tool (28), additional promoters were mapped with the `PlatProm` algorithm (29). The VirF binding sites were predicted manually based on phylogenetic footprinting of known binding regions. A+T tracks were classified as tracks if six or more A or T were present at the same time.

### Modeling and visualization of protein structures

The three-dimensional structures of the IpaH proteins from *Escherichia marmotae* were modeled using the `Swiss-Model` program (30) employing PDB: 5KH1.1 as the template. As a visualization tool, the UCSF `Chimera` software was used (31).

## Results

### Validation of genome assemblies

We analysed 130 *Shigella genomes* including 46 *S. flexneri,* 25 *S. dysenteriae,* 19 *S. boydii,* 39 *S. sonnei*, and one unclassified *Shigella* strain (*Supplementary Table 1*). We used two criteria to validate the *Shigella* annotation, the presence of the *ipaH* genes and other components of T3SS ***(Table 1)***. As the T3SS markers we used the *mxiC, mxiE, mxiG, virB, virF, spa15, spa32, spa40, ipgA, ipgB, ipgD, apaA, ipaB, ipaC, ipaD, mxiH, icsB* genes. For three assemblies, we have found neither *ipaH* nor T3SS hits. These samples were extracted from soil, stream sediment, and Antarctic lichen so we classified them as non-invasive *E. coli* and excluded them from the analysis. Additionally, we checked that non-invasive *E. coli* strains did not have any of these virulence determinants using the set of 414 *E. coli + Shigella* genomes from (32). In 17 assemblies, the plasmids were absent but we found chromosomal *ipaH* genes. 37 assemblies comprised plasmids but none of them held the components of T3SS. These results may be explained by elimination of the plasmids during cultivation (33). Only 64 assemblies contained all essential virulence elements.

**Table 1.**
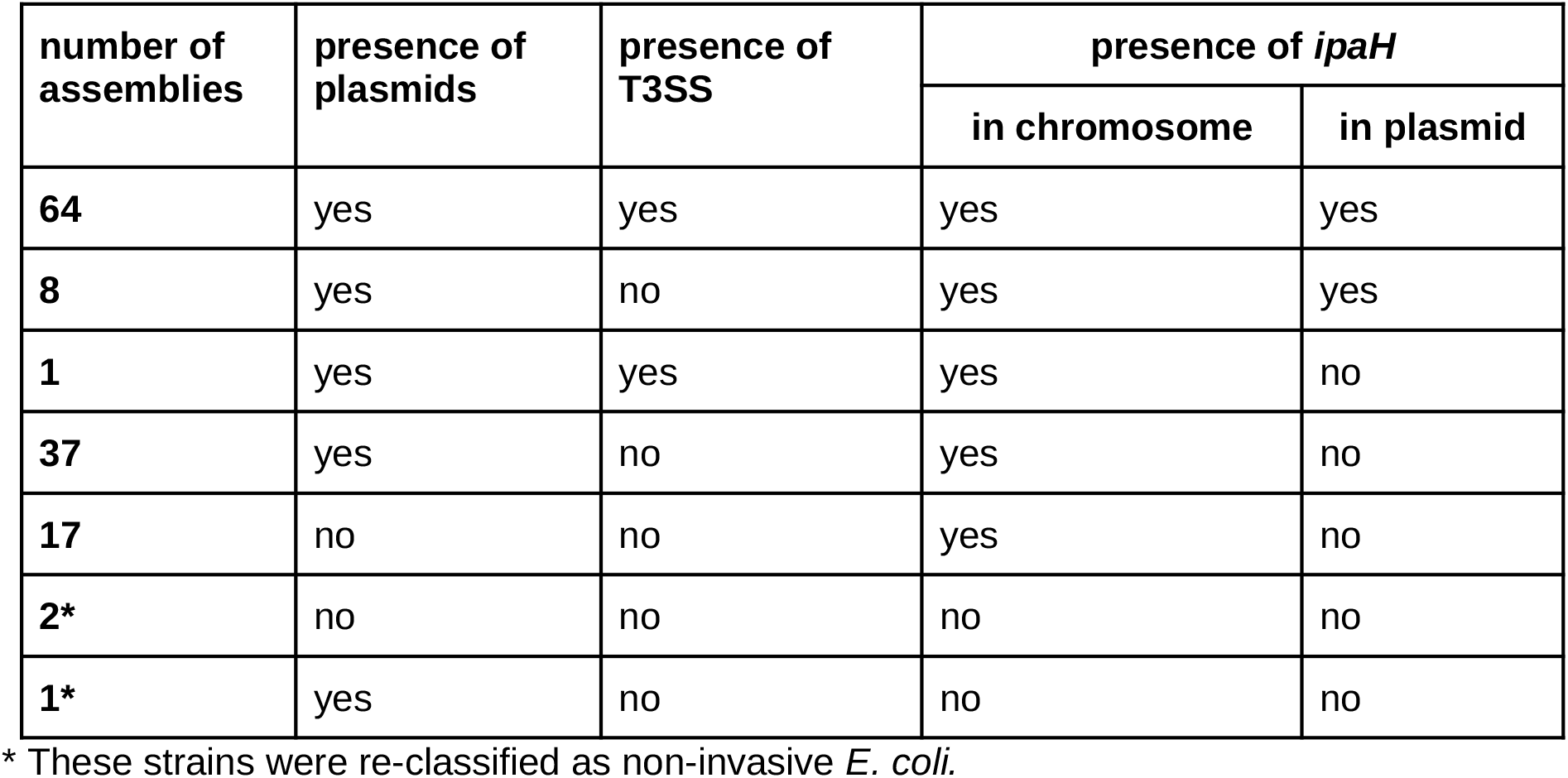
Statistics of *Shigella* assemblies.

We also characterised the *ipaH* genes in three available EIEC lineages from (1). One strain (*E. coli* NCTC 9031) did not contain *ipaH* genes or T3SS genes, thus the strain was filtered out. Two other strains (*E. coli* CFSAN029787 and *E. coli* 8-3-Ti3) had *pINV* with genes of the T3SS system, and the *ipaH* genes on the chromosomes and plasmids **(*Supplementary Table 1*)**.

### Classification of the *ipaH* genes

There is no consistent nomenclature of the *ipaH* genes across *Shigella* strains and the number of the *ipaH* genes in a strain varies [see Table 1 in (34)], thus, we created a unifying classification of all *ipaH* gene family members. In 127 *Shigella* assemblies, we found 864 protein sequences belonging to the E3 ubiquitin-ligase family (see Methods, ***Supplementary Table 3***). Based on sequence similarity of the recognition domains and the composition of regulatory elements in upstream regions, we divided all *ipaH* genes into nine classes ***(Figure 1)***. Confirming this classification, proteins from different classes were also distinguishable by their length, the number of LRRs, and the length of conserved upstream regions ***(Table 2)***. Taking into account high sequence similarity of genes across *Shigella*, we used consensus *ipaH* sequences (***Supplementary Table 5)*** from each class for gene annotation.

**Table 2.**
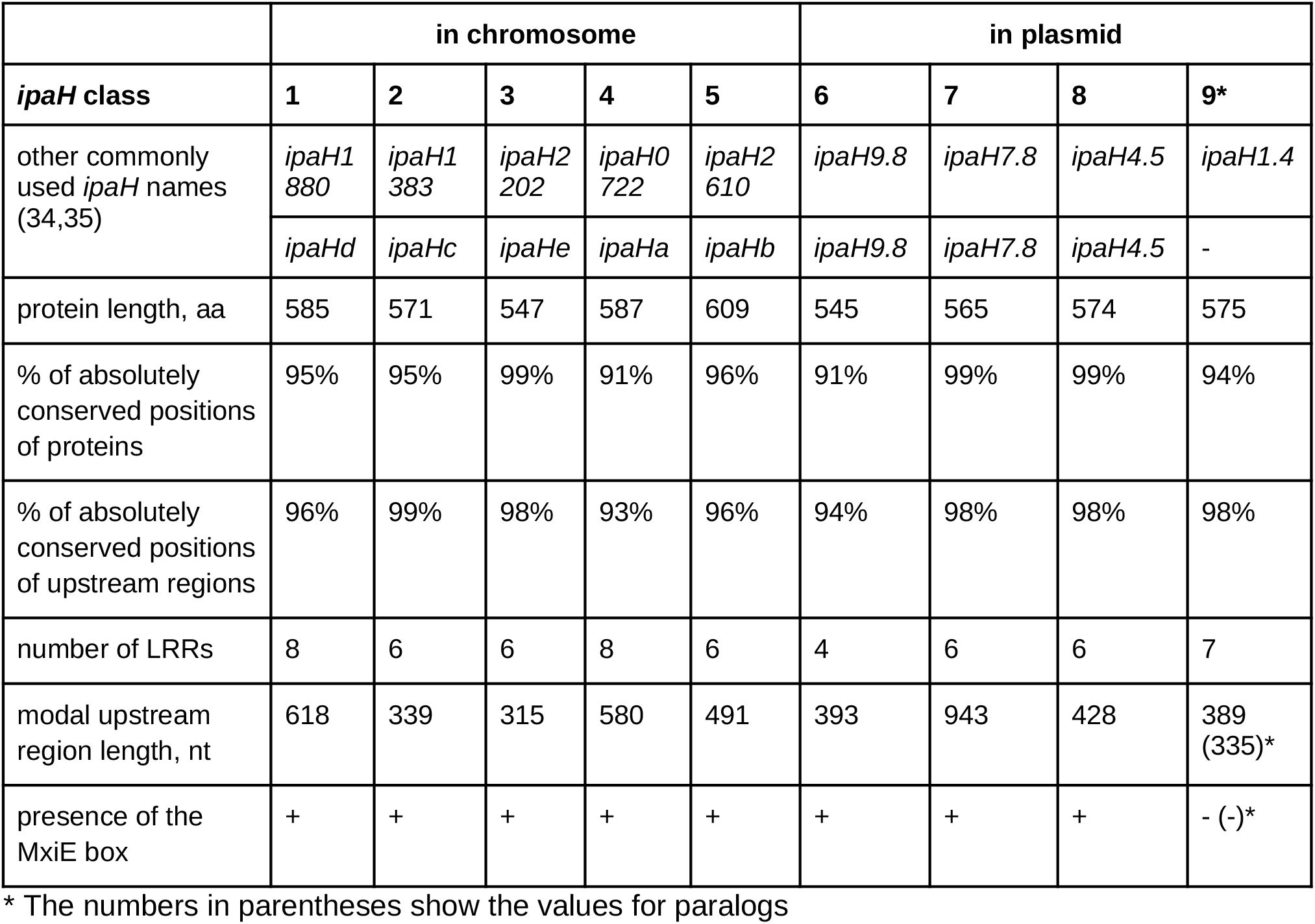
Classification of the *ipaH* genes from *Shigella*: coding sequences and upstream regions.

|  | in chromosome |  |  |  |  | in plasmid |  |  |  |
| --- | --- | --- | --- | --- | --- | --- | --- | --- | --- |
| *ipaH* class | 1 | 2 | 3 | 4 | 5 | 6 | 7 | 8 | 9* |
| other commonly used *ipaH* names (34,35) | *ipaH1*<br>880 | *ipaH1*<br>383 | *ipaH2*<br>202 | *ipaH0*<br>722 | *ipaH2*<br>610 | *ipaH9.8* | *ipaH7.8* | *ipaH4.5* | *ipaH1.4* |
|  | *ipaHd* | *ipaHc* | *ipaHe* | *ipaHa* | *ipaHb* | *ipaH9.8* | *ipaH7.8* | *ipaH4.5* | - |
| protein length, aa | 585 | 571 | 547 | 587 | 609 | 545 | 565 | 574 | 575 |
| % of absolutely conserved positions of proteins | 95% | 95% | 99% | 91% | 96% | 91% | 99% | 99% | 94% |
| % of absolutely conserved positions of upstream regions | 96% | 99% | 98% | 93% | 96% | 94% | 98% | 98% | 98% |
| number of LRRs | 8 | 6 | 6 | 8 | 6 | 4 | 6 | 6 | 7 |
| modal upstream region length, nt | 618 | 339 | 315 | 580 | 491 | 393 | 943 | 428 | 389 (335)* |
| presence of the MxiE box | + | + | + | + | + | + | + | + | - (-)* |

\* The numbers in parentheses show the values for paralogs

**Figure 1.**
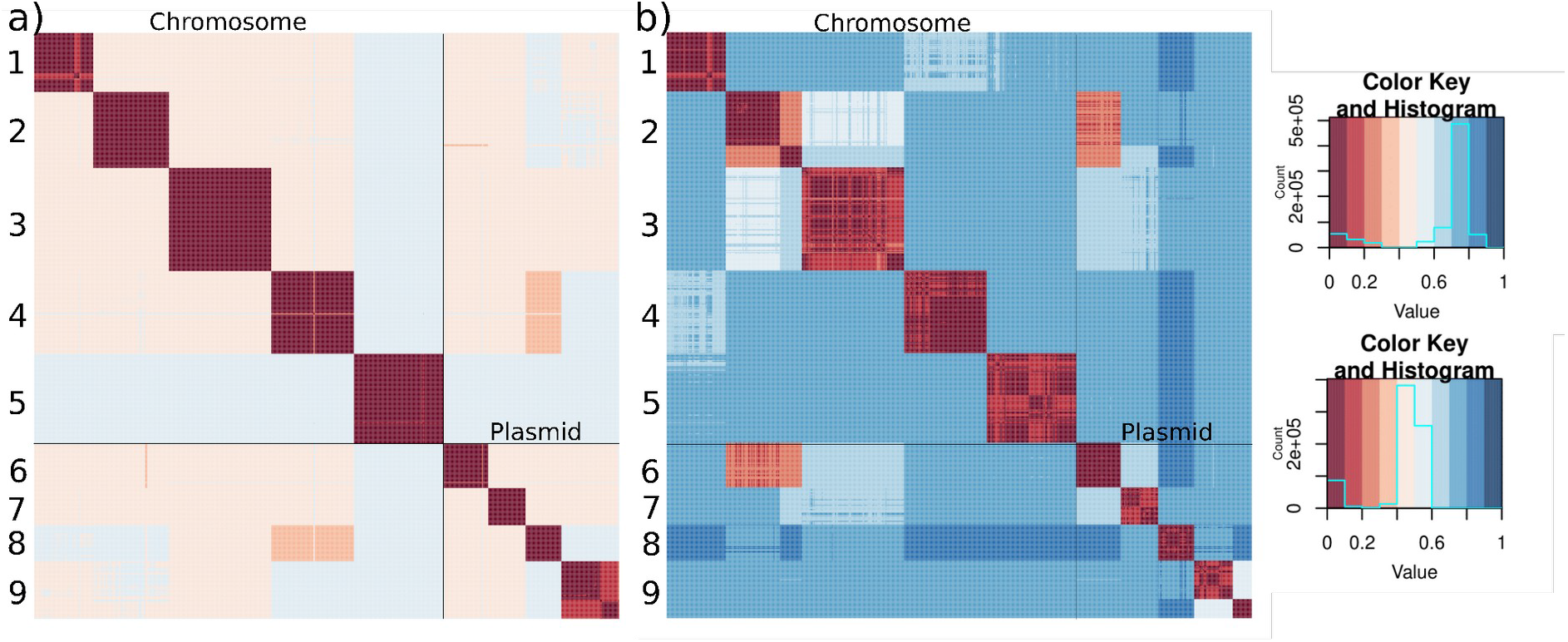
Heatmap of the identity levels of (a) the *ipaH* genes; (b) their upstream sequences *in Shigella*. Pairwise distances were calculated as 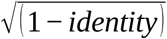.

Interestingly, the *ipaH* genes from classes #1-5 were present only in chromosomes while those from classes #6-9 were found only in plasmids. The only exception was a duplicated *ipaH* gene from class 5 in *Shigella flexneri* 1a strain 0228, where one copy was encoded in the chromosome and the other one in the plasmid. This assembly did not contain the *pINV* plasmid with T3SS genes, thus the observation might have been caused by miss-assembly. Genes from classes #4 and #8 had the highest protein sequence similarity while upstream regions were most similar for genes from classes #2 and #6.

Only 45% of the *Shigella* genomes hold a complete set of chromosomal *ipaH* genes and 20% genomes have a complete set of plasmid *ipaH* genes (for plasmids this is a lower-bound estimate as many assemblies lack plasmid sequences). Moreover, in many genomes *ipaH* classes #3, #5, and #9 were represented by more than one copy. Most *ipaH* copies were identical, the exception is two subclasses (#9a and #9b) that were distinguishable both by their gene and upstream sequences ***(Figure 1)***. Subclass #9b was found in almost all *Shigella flexneri* genomes, so we hypothesized that the *ipaH* #9b copy had been acquired by the common ancestor of the S. *flexneri* branch.

### Regulatory patterns in the *ipaH* upstream regions

In addition to the high level of sequence similarity in each *ipaH* class, the upstream regions of the genes were also highly conserved. Indeed, the upstream intergenic regions of different *ipaH* genes comprised 300-900 base pairs with identity of more than 90% in each class, except class #9 (see below). Interestingly, the similarity was high starting from the translation start codon to (and including) putative binding sites of transcription factor MxiE, especially in classes #2 and #6, suggesting a key role of MxiE in the regulation of *ipaH* transcription. Previously, the relative positioning of MxiE binding sites and transcription starts, as well as sequences of the MxiE box, -10 box, and the spacer between them were used to classify the *ipaH* genes into eight regulatory classes (34). Each class defined by our sequence similarity approach, except for class #9, corresponds to one of the regulatory classes ***(Figure 2B)***. Indeed, each class has its unique regulatory pattern characterized not only by the MxiE-box positioning and the spacer sequence, but also by the presence of A+T rich tracks as possible targets for the interaction with VirF and H-NS. Specifically, classes #4, #5, and #7 possess both A- and T-rich tracks (see ***Figure 2A*** for an example), classes #1 and #2 has mainly polyT-tracks, and class #3 has mainly polyA-tracks.

**Figure 2.**
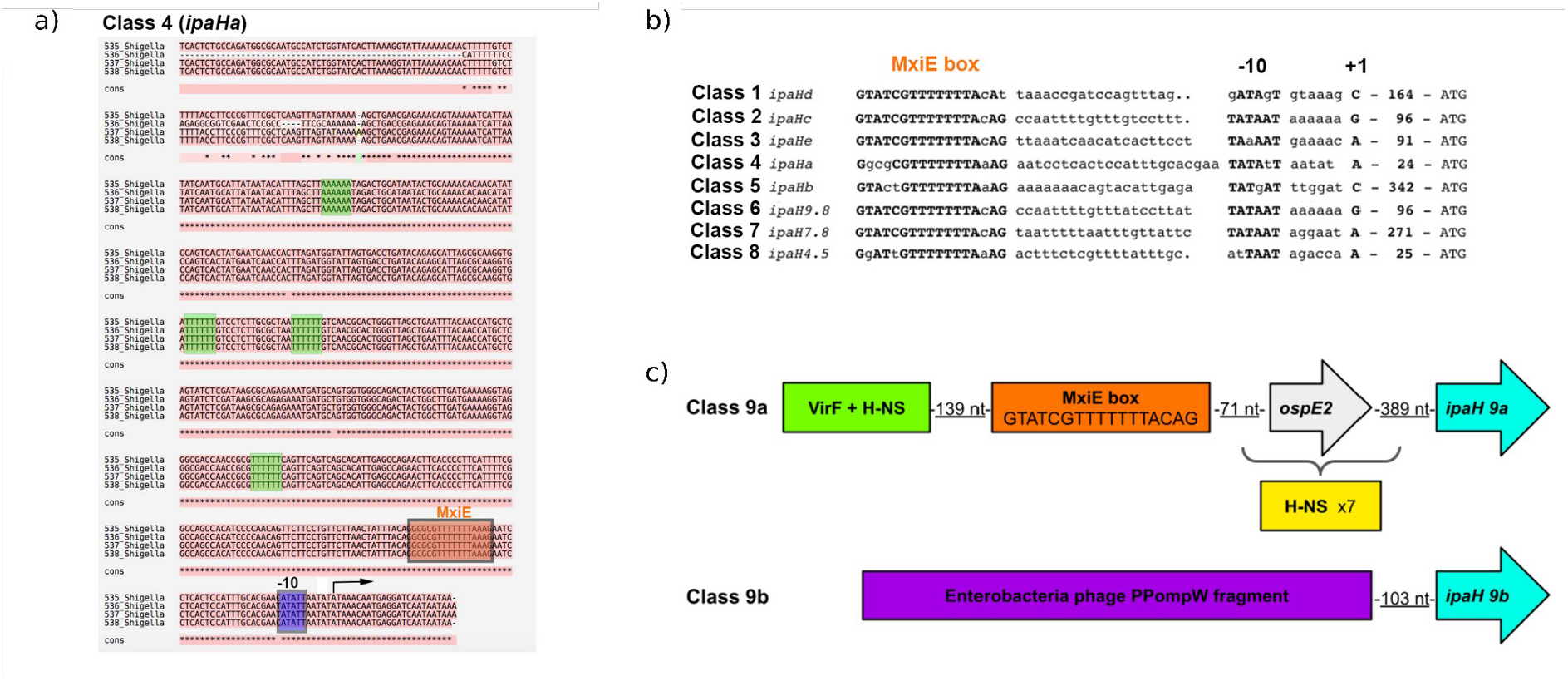
Regulatory elements in *ipaH* upstream regions. a) Alignment of the upstream region for selected representatives of the *ipaH* class #4. The putative MxiE box is indicated by the orange box, A/T tracks are in green boxes. The transcription start is indicated by the black arrow. b) Comparison of the sequence-based *ipaH* classification with the classification based on positioning of the MxiE box, -10 element, and the sequence of the spacer. Adapted from (34). c) Principal scheme of upstream regions of the *ipaH* classes #9a and #9b.

Plasmid *ipaH* genes of class #9 were divided into two groups. Genes from class #9b had disrupted upstream regions due to a prophage insertion and thus did not appear to have regulatory elements typical for other *ipaH* classes ***(Figure 2C)***. Also, no candidate promoters upstream of *ipaH* class #9b could be identified, suggesting that these genes may not be transcribed. The genes of class #9a also did not have an upstream MxiE box, however, they might be transcribed polycystronically with the *ospE* gene ***(Figure 2C)*** utilizing its regulatory elements. The *ipaH* genes from class #9a were surrounded by multiple A+T-rich tracks typical for mobile elements or prophages (17).

The upstream regions of different *ipaH* classes are not similar at any significant level, the only exceptions being classes #2 (chromosomal) and #6 (plasmid) that have a highly similar 150 bp fragment of the regulatory region between the MxiE box and the translation start codon ***(Figure 2B)***.

### Phyletic patterns of *ipaH*

We analyzed the phyletic patterns of *ipaH* in *Shigella* and EIEC strains **(*Figure 3, Supplementary Table 3, Supplementary figure 1*)**. The reconstructed phylogenetic tree was generally consistent with previous reconstructions (1) and revealed five major *Shigella* clades with the tree topology not reflecting the species names. In our dataset, *S. sonnei* and *S. flexneri* were monophyletic (marked in *yellow* and *violet* in **Figure 3**, respectively), *S. boydii* and *S. dysenteriae* were mixed in two distant clades (marked in *orange* and *red* in **Figure 3**, respectively) and a set of *S. dysenteriae* strains formed the fifth clade (the *green* clade in **Figure 3**). The phyletic patterns of the *ipaH* genes were highly mosaic. Nevertheless, we observed some clade-specific patterns. In particular, class #1 was rare in the *orange* clade, while class #3 was absent in the *green* clade.

**Figure 3.**
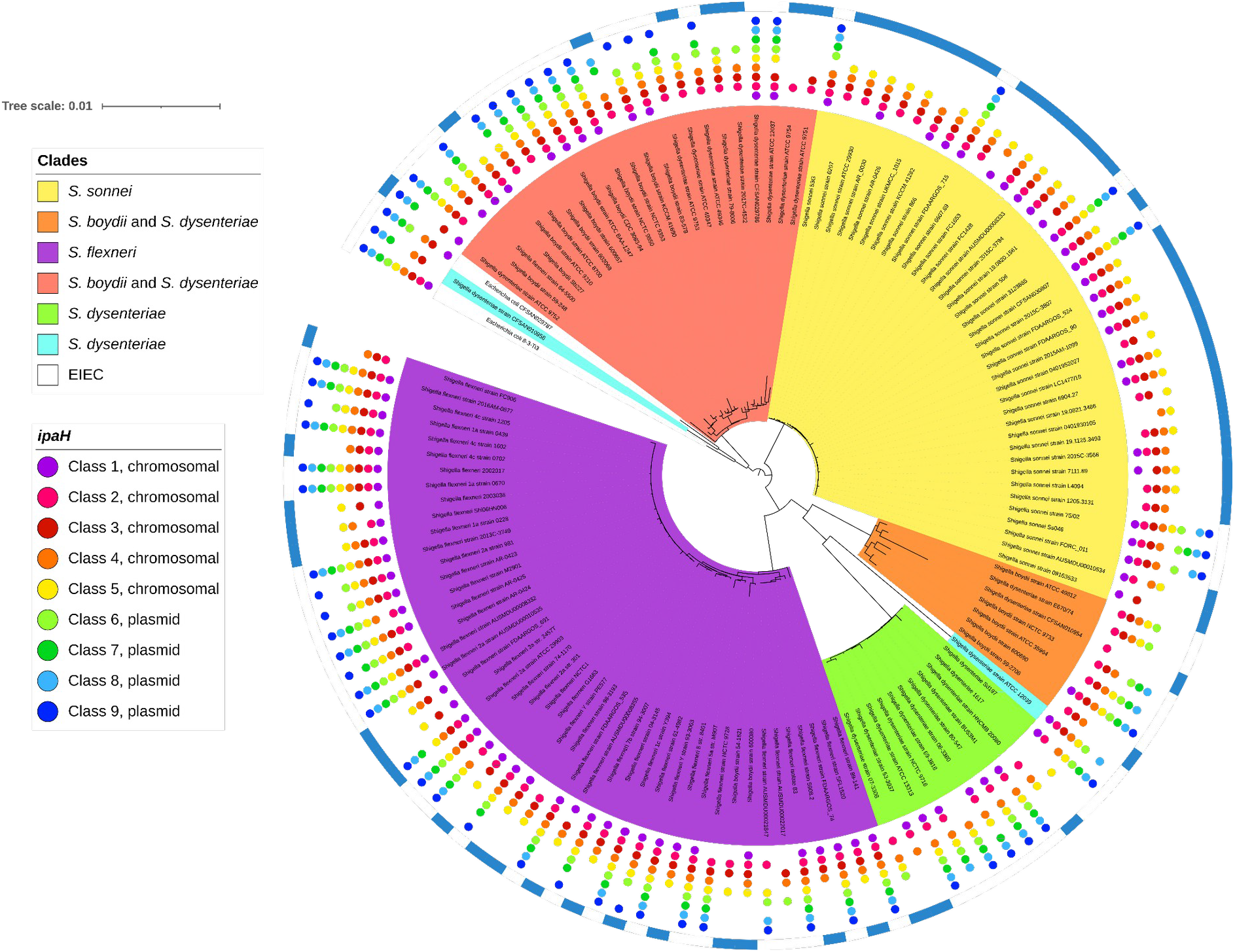
Phyletic patterns of the *ipaH* genes in *Shigella* and EIEC. The coloring of the unrooted tree reflects major *Shigella* clades that putatively evolved from different non-pathogenic *E. coli;* two distant *Shigella* strains are shown in blue, the EIEC strains are shown in white. The presence of the *ipaH* genes is shown by dots whose color reflects the *ipaH* class (see the legend). The genes in classes #1-5 are located in chromosomes; the genes in classes #6-9, in plasmids. The genomes marked by the external blue arcs do not contain the T3SS genes.

The strains of EIEC did not cluster with the major *Shigella* clades or with each other (**Figure 3**). The gene content and their genomic distribution was also consistent with polyphyletic origin of the EIEC strains. Specifically, *Escherichia coli* 8-3-Ti3 had a complete set of *ipaH*, while in *Escherichia coli* CFSAN029787, two chromosomal *ipaH* genes were missing. These genes were not distinguishable from *Shigella* effectors, and their location on chromosomes and plasmids was consistent with their class assignments. In *Escherichia coli* CFSAN029787, *ipaH* #1 and *ipaH* #3 had frameshifts, likely resulting in pseudogenization.

Interestingly, copies of *ipaH* genes were found in many *Shigella* genomes both on chromosomes and plasmids **(*Supplementary Table 3)***. We observed paralogs of *ipaH* #2, #4, #5 in the *orange (boydii & dysenteriae)* clade, only *ipaH* #4 in the *green (dysenteriae)* clade, and only *ipaH* #3 in the *red (boydii & dysenteriae)* clade **(*Supplementary Figure 2a*)**. Genomes of the *violet (flexneri)* clade had paralogs of the *ipaH* #3, #4, #5, #7, and #9 **(*Supplementary Figure 2b*)**, while genomes in the *yellow* (*sonnei*) clade did not have *ipaH* duplicates **(*Supplementary Figure 2c*)**. Surprisingly, none of *ipaH* paralogs were tandem repeats; in contrast, the copies located at some distance from each other and frequently surrounded by prophages and pseudogenes.

### *ipaH* repertoire in non-human *Escherichia*

We identified and compared the *ipaH* genes in pathogenic *Escherichia* spp. extracted from non-human hosts ***(Supplementary Table 4, Figure 4)***. Previously, nine *ipaH* genes and two short ORFs containing fragments of *ipaH* genes were found in the genome of *Escherichia marmotae* HT073016, isolated from faecal samples of *Marmota himalayana* (20). The authors reported automated annotation of eleven genes as *ipaH*: four on the *pEM148* plasmid, five on the *pEM76* plasmid, and two on the chromosome. According to our *ipaH* identification procedure (see Methods) we confirmed eight of these gene annotations and found one additional chromosomal gene. We excluded from the analysis short ORFs that did not contain the N-terminal domain assuming these to be mis-annotation or remains of pseudogenes.

**Figure 4.**
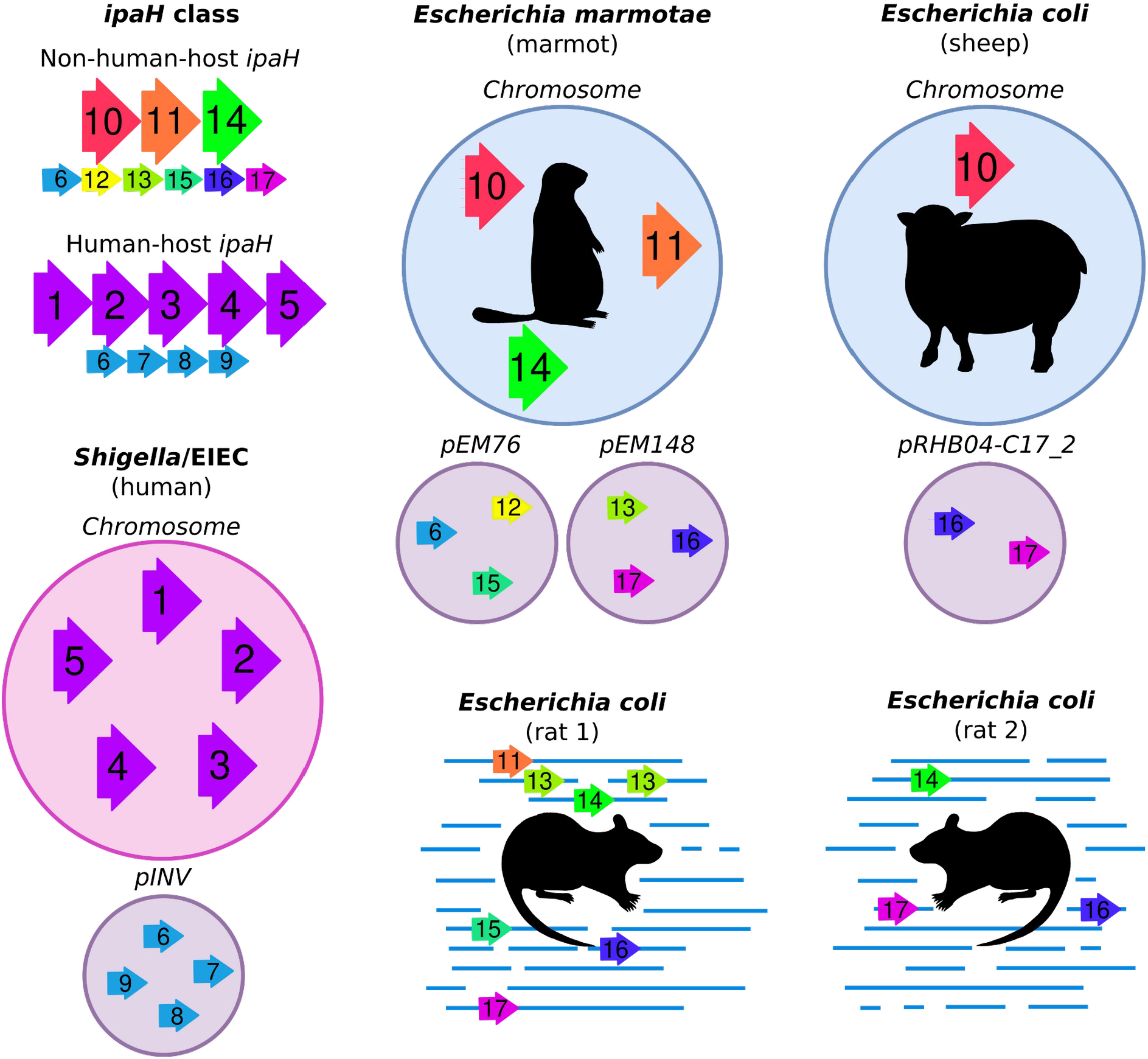
Composition of the *ipaH* genes in the *Escherichia* genomes from different hosts. Assemblies of marmot- and sheep-host *Escherichia* are complete, genomes of rat-host *Escherichia* are assembled as contigs. For *Shigella*, a genome with a complete set of the *ipaH* genes is shown.

In addition, we observed that *Escherichia coli* extracted from non-human hosts contained T3SS as well as *ipaH* genes. Specifically, two strains extracted from rat feces, *Escherichia coli* CFSAN092688 and *Escherichia coli* CFSAN085900 had six and three *ipaH* genes, respectively, and a strain from pooled sheep faecal samples, *Escherichia coli* RHB04-C17, had three *ipaH* genes.

Based on sequence similarity of recognition domains, we classified the IpaH proteins from non-human-host *E. coli* into nine classes ***(Figure 5a)***. The level of sequence similarity between non-human hosts IpaH that belong to the same class is less than that for *Shigella*. Based on the location of the *ipaH* genes in completely assembled genomes of marmot- and sheep-host *Escherichia*, we assume that they retain their location in replicons.

**Figure 5.**
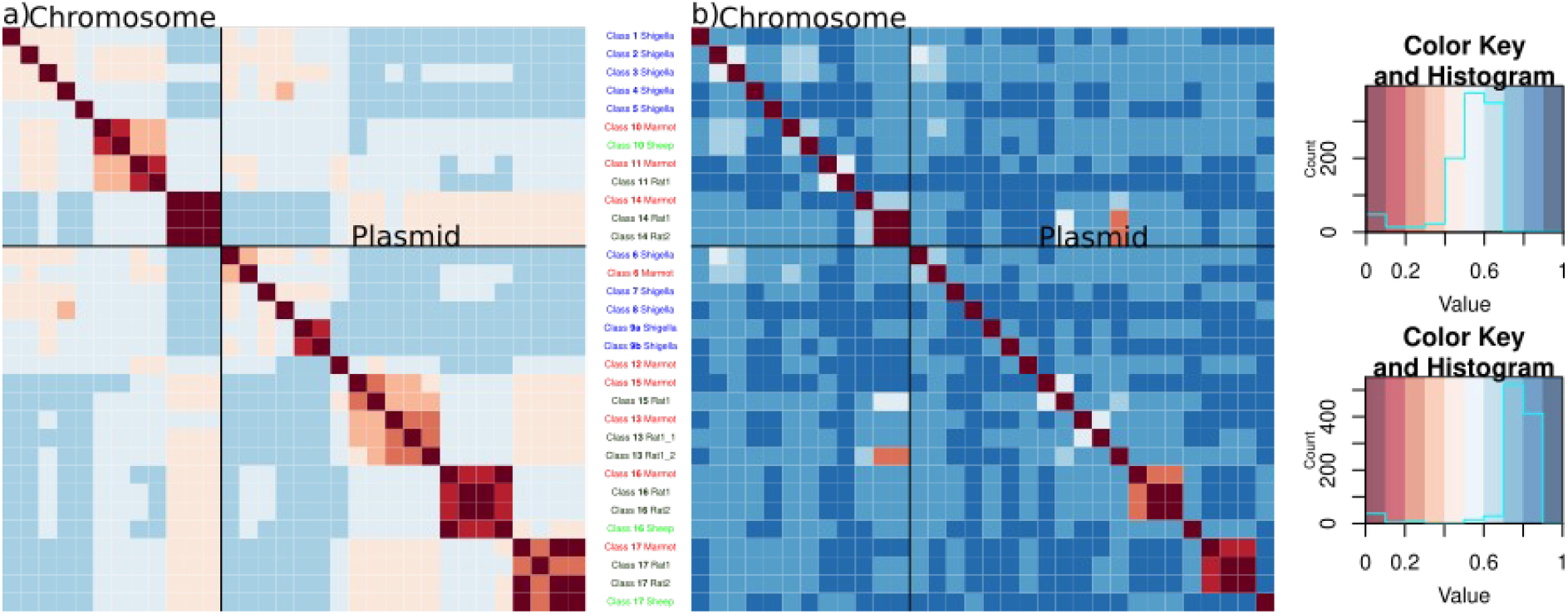
Heatmap of the identity level of (a) the *ipaH* genes; (b) their upstream sequences in non-human-host *Escherichia* spp. Hosts are labeled according to the following principle: marmot is marked by red, rat is marked by dark green, sheep is marked by light green. Representative sequences of *Shigella ipaH* genes also were included in comparison, their labels marked in blue. Pairwise distances were calculated as 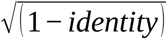.

Two *ipaH* classes #16 and #17, putatively in plasmids, were found in all non-human-host *Escherichia* spp. Putatively chromosomal *ipaH* class #14 were present in marmot-host and rat-host *Escherichia* spp.; in turn, the genomes of marmot- and sheep-host *Escherichia* spp. share *ipaH* class #10. Only one of non-human-host *Escherichia ipaH* genes (class #6) was present in *Shigella*, however the upstream sequences of *ipaH* class #6 in *Shigella* and *Escherichia marmotae* were significantly different ***(Figure 5b)***. In particular, the regulatory regions of *ipaH* from non-human-host *Escherichia* spp. contain neither MxiE boxes, nor multiple A/T tracks.

The upstream regions of most *ipaH* classes were similar in rat-host and marmot-host *E. coli*, while the upstream regions in sheep-host *E. coli* genes and all *ipaH* upstream regions were unique. Classes #13 (in plasmid) and #14 (in chromosome) show gene-sequence and upstream-sequence similarity but have different numbers of LLRs, which indicates their evolution by gene duplication and subsequent deletions or tandem duplication of short genomic segments.

Surprisingly, in non-human hosts *Escherichia* spp., the C-terminal domain of IpaH proteins was not conserved ***(Supplementary Table 6)***. Non-human hosts IpaH classes #10, #11, #12 had a C-terminal domain similar to that in *Shigella* (92% aa identity), while the C-terminal domain of IpaH classes #13 through #17 was more diverged (75% aa identity) ***(Figure 6)***. This observation also explains the results from (20) that only a fraction of the identified *ipaH* genes were homologous to the *ipaH* of *Shigella* spp.

**Figure 6.**
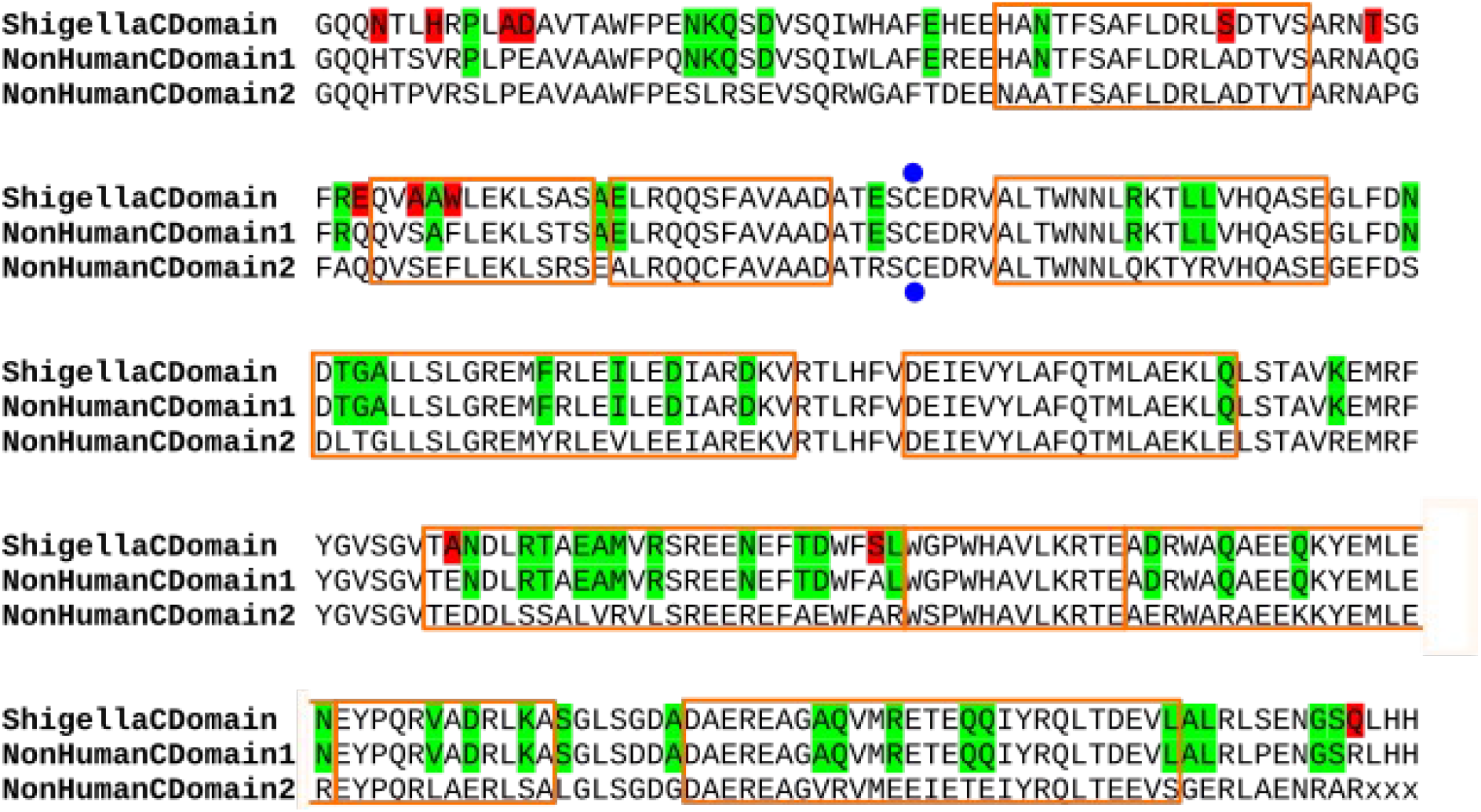
**Alignment of the IpaH C-terminal domains** from *Shigella* spp. and *Escherichia marmotae*. Consensus sequences are shown. The active site is marked in blue dots, alpha-helices are shown by orange frames. The differences between *Shigella* and both *E. marmotae* proteins are marked in red, the differences between two types of the C-terminal domains in *E. marmotae* are marked in green.

We mapped the amino acid substitutions between consensus sequences of C-terminal domains of IpaH from *Shigella* spp. and non-human-host *Escherichia* spp. on the three-dimensional structure of *Shigella flexneri* effector IpaH1880 (PDB: 5KH1) **(*Figure 6, Supplementary Figure 3*)**. These differences were not clustered, nor did they affect the protein active site.

## Discussion

*Shigella* spp. and enteroinvasive *E. coli* have a wide variety of IpaH effectors that play a significant role in invasion, modulation of inflammation, and host response (3). Previously, several studies attempted to describe the *ipaH* gene family in *Shigella* aimed to compare representative strains from different *Shigella* species (34,35). However, *Shigella* spp., and the known EIEC lineages, are paraphyletic with highly variable genomes. Therefore, a comprehensive comparative analysis combining all available genomic data was required to obtain a general picture of the gene family composition and evolution.

Collecting a large set of *ipaH* genes, we classified them based on sequence similarity and unified their nomenclature while maintaining references to previously used gene names (34,35). Although the sequences of most *ipaH* genes were highly conserved across strains, in class #9 (*ipaH1.4*) we detected paralog diversification that might indicate formation of a new *ipaH* class. Given the important role of this family in the *Shigella* virulence, a consistent gene annotation is of direct medical relevance. Our results suggest that using consensus *ipaH* sequences from each class for gene annotation is efficient, reduces errors in annotation, and might be useful for future studies of this gene family.

The presence of IpaH effectors is one of the markers used for *Shigella* serotyping (36), however, less than a half of sequenced genomes had the entire set of the *ipaH* genes. Contrary to previous observations on smaller datasets (32), none of the *ipaH* genes are common to all *Shigella* strains, and the phyletic patterns of the *ipaH* genes suggest numerous independent gene losses. While the targets of some IpaH proteins are unknown, some proteins have shown to affect the same pathway at different stages, working together to cause disease (3). In this case, a complete set of IpaH would be functionally redundant and may not necessarily be preserved. Note that in case of bacterial isolates, elimination of plasmids and virulence factors in the course of cultivation may have led to the loss of plasmid classes prior to genome sequencing (33).

The presence of non-tandem copies of *ipaH* genes with conserved upstream regions in many *Shigella* strains indicate the acquisition of DNA fragments with *ipaH* from the same source and functionality and specificity of *ipaH* upstream regions. Superficially, the chromosomal *ipaH* genes seem to have more A+T tracks and presumably more options for regulation than those located on the plasmid. Indeed, the virulence plasmids likely have resulted from multiple events of transmission and transposition, and may only hold elements absolutely necessary for fast switches between cell functional states.

We did not detect any consistent differences in the repertoire of the *ipaH* genes in the *Shigella* and EIEC pathotypes. Moreover, the regulatory patterns in the upstream regions were the same. As notation of *Shigella* and EIEC pathotypes is not strongly defined, it is still not clear whether these factors are responsible for the differences in the infectious dose and disease severity of *Shigella*/EIEC pathotypes.

Interestingly, the *ipaH* composition and regulatory patterns in non-human host derived *Escherichia* differed substantially from the human host-derived strains. In total we detected eight new classes of the IpaH effectors in non-human host *Escherichia* spp. As the human-host *Escherichia* spp., they maintain their location in the chromosome or plasmids. One *ipaH* class (#6, *ipaH9.8*) was present both in *Shigella* spp. and *Escherichia marmotae* plasmids, which indicates the possibility of horizontal gene transfer between *E. coli* adapted to different hosts. In contrast to *Shigella ipaH*, the regulatory regions of *ipaH* from non-human-host *Escherichia* spp., while featuring A/T-rich elements characteristic to promoter regions, contain neither MxiE boxes, nor multiple A/T tracks: the only example with two such tracks is *ipaH* class #16 from marmot and rat.

Surprisingly, in the IpaH proteins encoded in the *Escherichia* genomes from non-human hosts we found two diverse C-terminal domains. This observation may be explained by the acquisition of effectors horizontally as well as differentiation of their functional roles in non-human-hosts *Escherichia*.

The IpaH proteins are considered as a candidate target of antibiotics due to their *Shigella* specificity. The first strategy is to target the C-terminal domain as it is highly conserved among *Shigella* IpaH effectors (37). However, IpaH can affect the antimicrobial activity of host proteins even in the absence of catalytic activity (38). Thus, targeting N-domains may be more effective but this strategy requires understanding of the *ipaH* repertoire in specific strains. To date, approaches used for testing the presence of *Shigella* virulence factors do not distinguish the members of the IpaH family (39), thus development of gene specific primers is required.

The *ipaH* variants of invasive *Escherichia* spp. from wildlife and domestic animals will require additional study as they may contribute to human-pathogen evolution. Notably, the annotation of the source of *E. coli* samples may be misleading. In particular, the *E.coli* genome extracted from a sample of sheep feces collected from the farm floor (BioSample: SAMN15147991) might be contaminated by bacteria from another host, such as a rat living on a farm. If this were the case, the new variant of the C-terminal domain of IpaH proteins may be rodent-specific. Extensive sampling and subsequent genomic sequencing of *Escherichia* spp. from different hosts will shed light on the specificity of the invasion system and IpaH effectors to the pathogens’ hosts.

## Supporting information

All figures in good resolution

## List of abbreviations

T3SS: type 3 secretion system
*pINV*: plasmid of invasion
NEL: novel E3-ubiquitin ligase
LRR: leucine rich repeat
EIEC: enteroinvasive *Escherichia coli*
ORF: open reading frame

## Author statements

### Funding

This study was supported by the Russian Foundation for Basic Research (RFBR), Grant # 20-54-14005 and Fonds zur Förderung der wissenschaftlichen Forschung (FWF), Grant # I5127-B. The work of OB is supported by the European Union’s Horizon 2020 Research and Innovation Programme under the Marie Skłodowska-Curie Grant Agreement No. 754411. The funders had no role in study design, data collection and analysis, decision to publish, or preparation of the manuscript.

### Authors contribution

OOB conceived and designed the study. NOD and MNT analysed the data. MSG and FAK aided in interpreting the results. All authors wrote, read, and approved the final version of the manuscript.

### Conflicts of interest

The authors declare that they have no competing interests.

### Ethical approval

Not applicable.

### Consent for publication

Not applicable.

## Acknowledgements

The project was initiated with Aygul Minnegalieva and Yulia Yakovleva at the Summer School of Molecular and Theoretical Biology (SMTB-2020), supported by the Zimin Foundation. We thank Inna Shapovalenko, Daria Abuzova, Elizaveta Kaminskaya, and Dmitriy Zvezdin for their contribution to the project during SMTB-2020. We also thank Peter Vlasov for fruitful discussions.

## Supplementary Figures

**Supplementary Figure 1.**
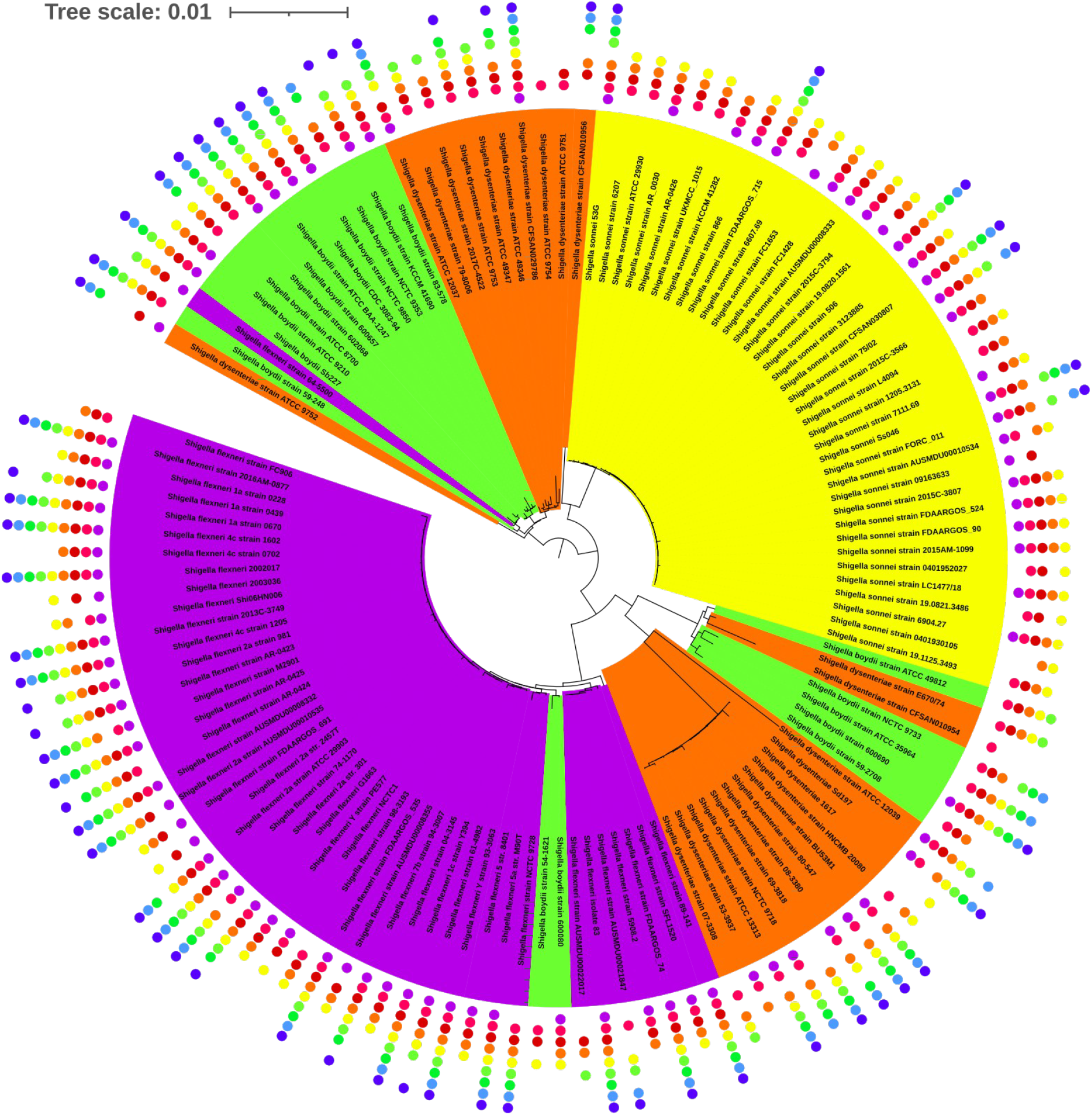
Phylogenetic tree of *Shigella* spp. colored by species name: violet – *S. flexneri*, green – *S. boydii*, orange – *S. dysenteriae*, yellow – *S. sonnei*. Other notation as in ***Figure 3***.

**Supplementary Figure 2.**
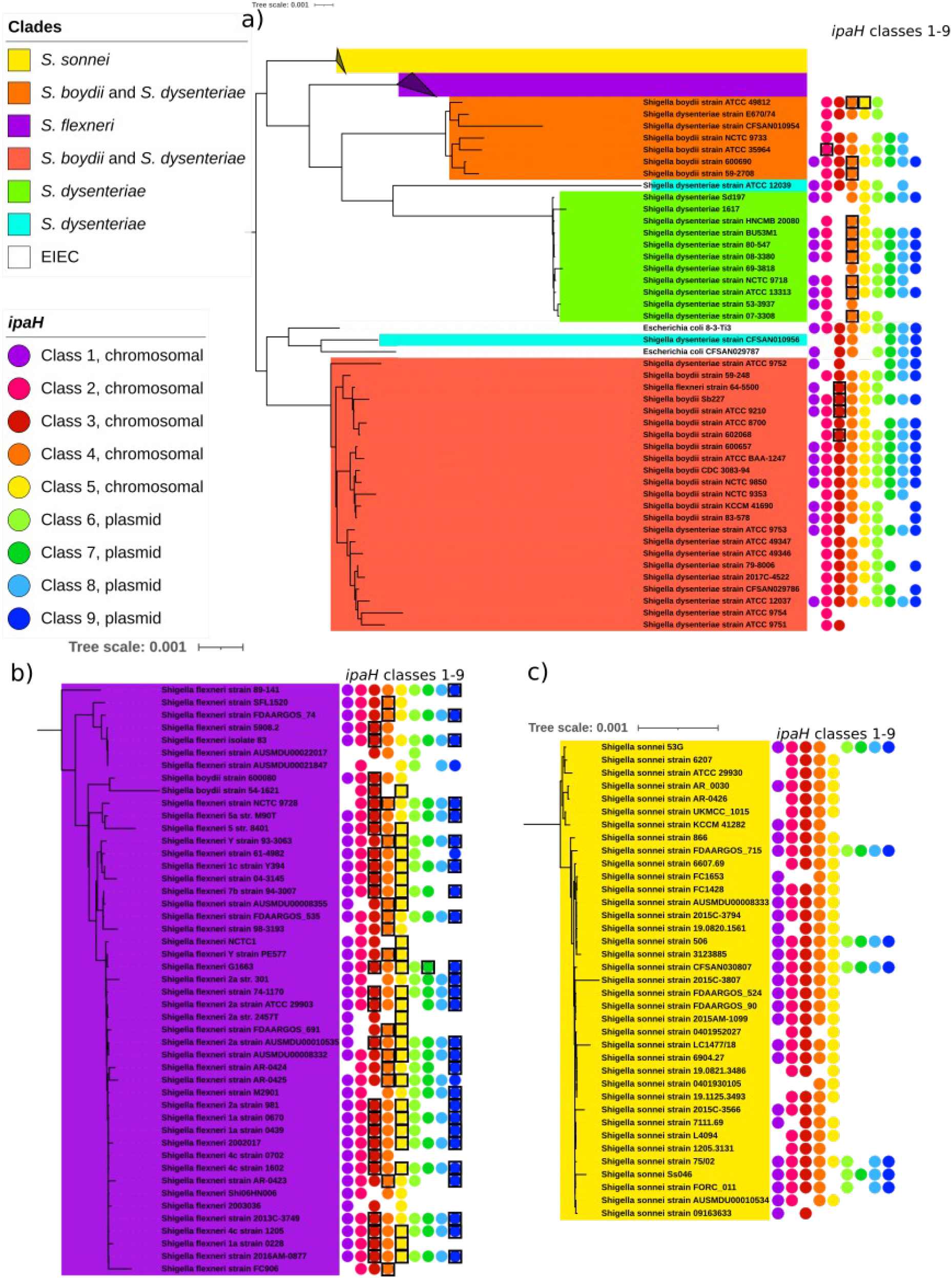
Abundance of the *ipaH* family members in *Shigella* clades. The notation as in ***Figure 3***. Black frames indicate duplications.

**Supplementary Figure 3.**
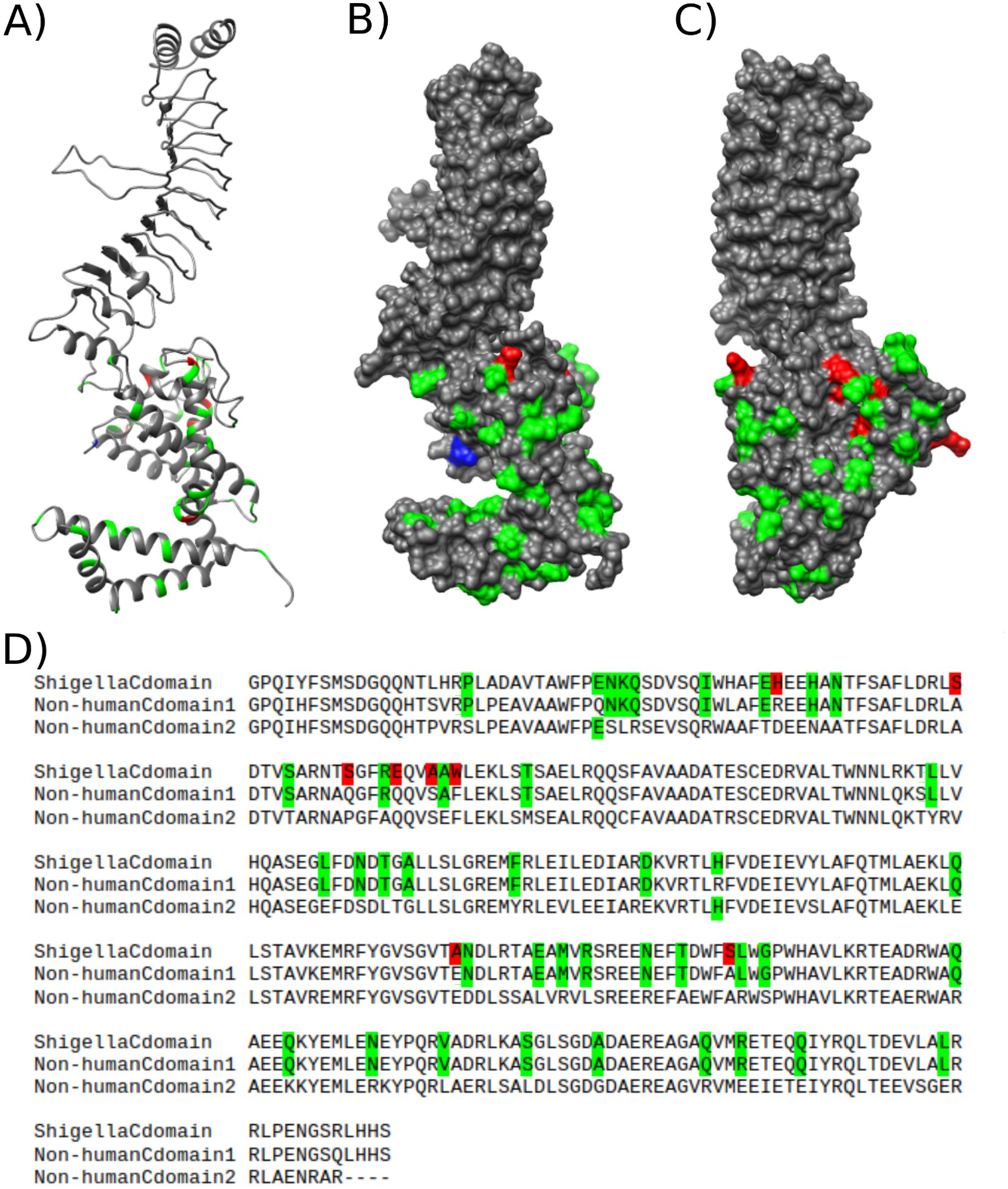
Homology modeling of IpaH from *Escherichia marmotae.* The predicted protein structure of *Escherichia marmotae* IpaH class 1 (A) as a ribbon representation; (B) with surface of the protein as seen from the binding site side; (C) with surface of the protein from the side opposite to the binding site. (D) Sequence alignment of three variants of the C-terminal domains from *Shigella* spp. and *Escherichia marmotae*. The active site is marked in blue. The positions strongly shared between two domains from *E. marmotae* but diverse in *Shigella* are marked in red; the positions strongly shared between *Shigella* and *Shigella*-type domain from *E. marmotae* are marked in green.

**Supplementary Table S1** Genomes of human-host invasive *Escherichia* and *Shigella*

**Supplementary Table S2** Genomes of non-human-host invasive *Escherichia*

**Supplementary Table S3** i*paH* genes from human-host invasive *Escherichia* and *Shigella*

**Supplementary Table S4** *ipaH* genes from non-human-host invasive *Escherichia* and *Shigella*

**Supplementary Table S5** Consensus sequences of IpaH proteins from human-host invasive *Escherichia* and *Shigella*

## Notes

**Data summary:** The datasets supporting the conclusions of this article are available at https://github.com/zaryanichka/E3UbLigases

### Competing Interest Statement

The authors have declared no competing interest.

