## Supplementary figures and images for "Chromosome-encoded IpaH ubiquitin ligases indicate non-human pathogenic *Escherichia*"

### Figure1.png

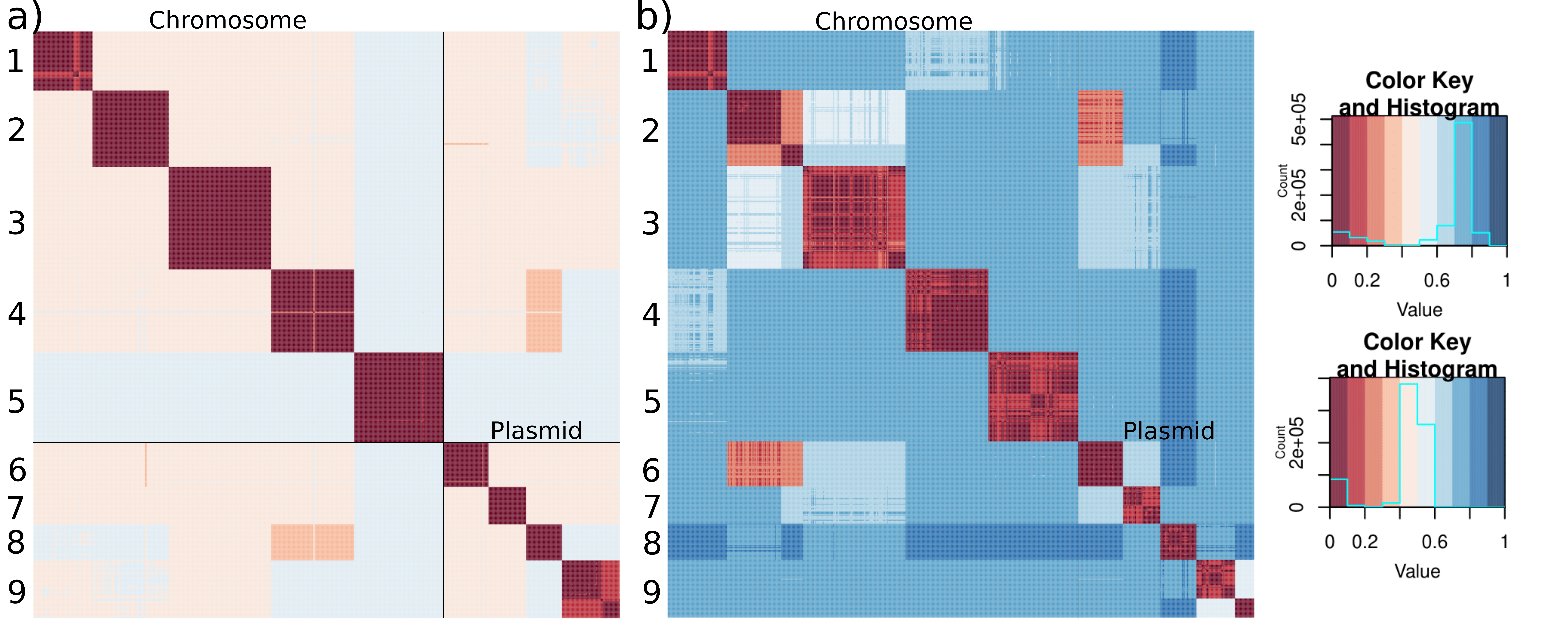

### Figure2.png

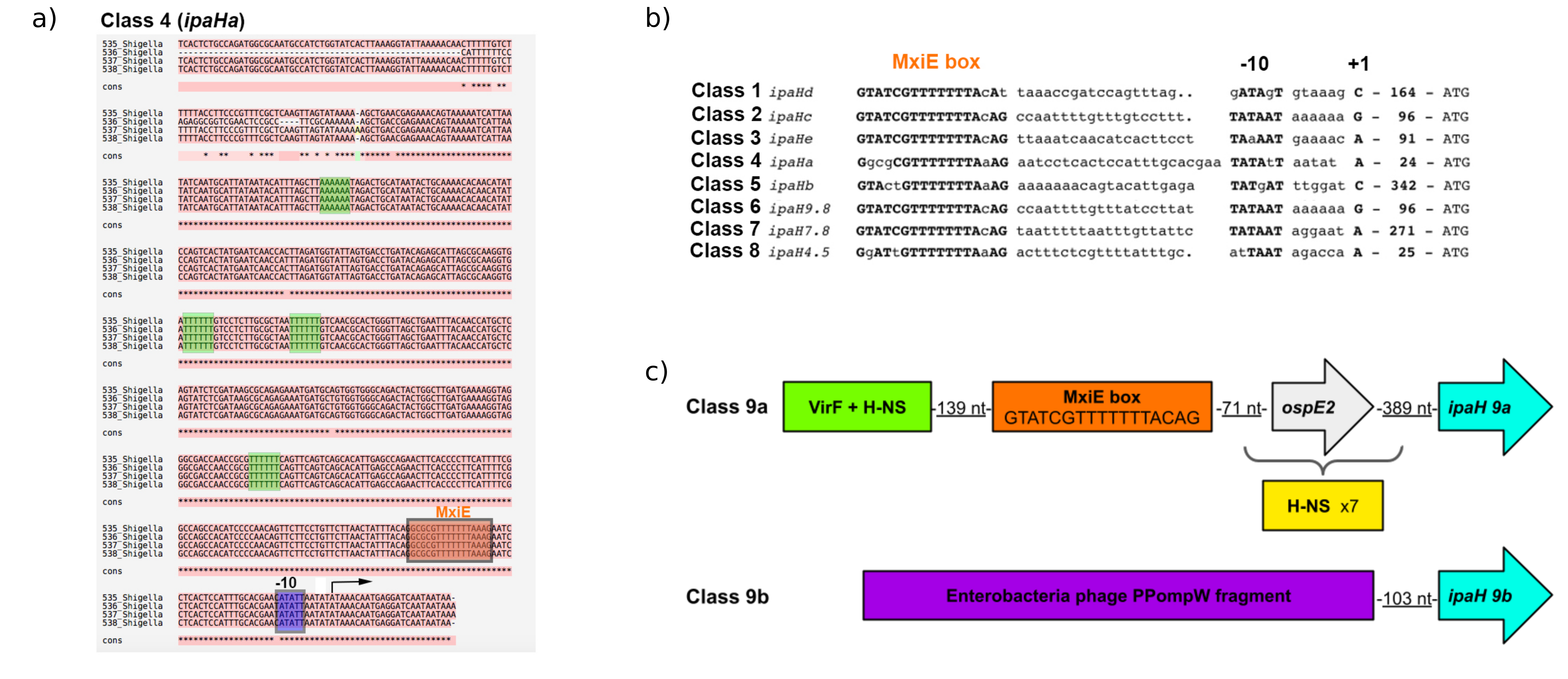

### Figure3.png

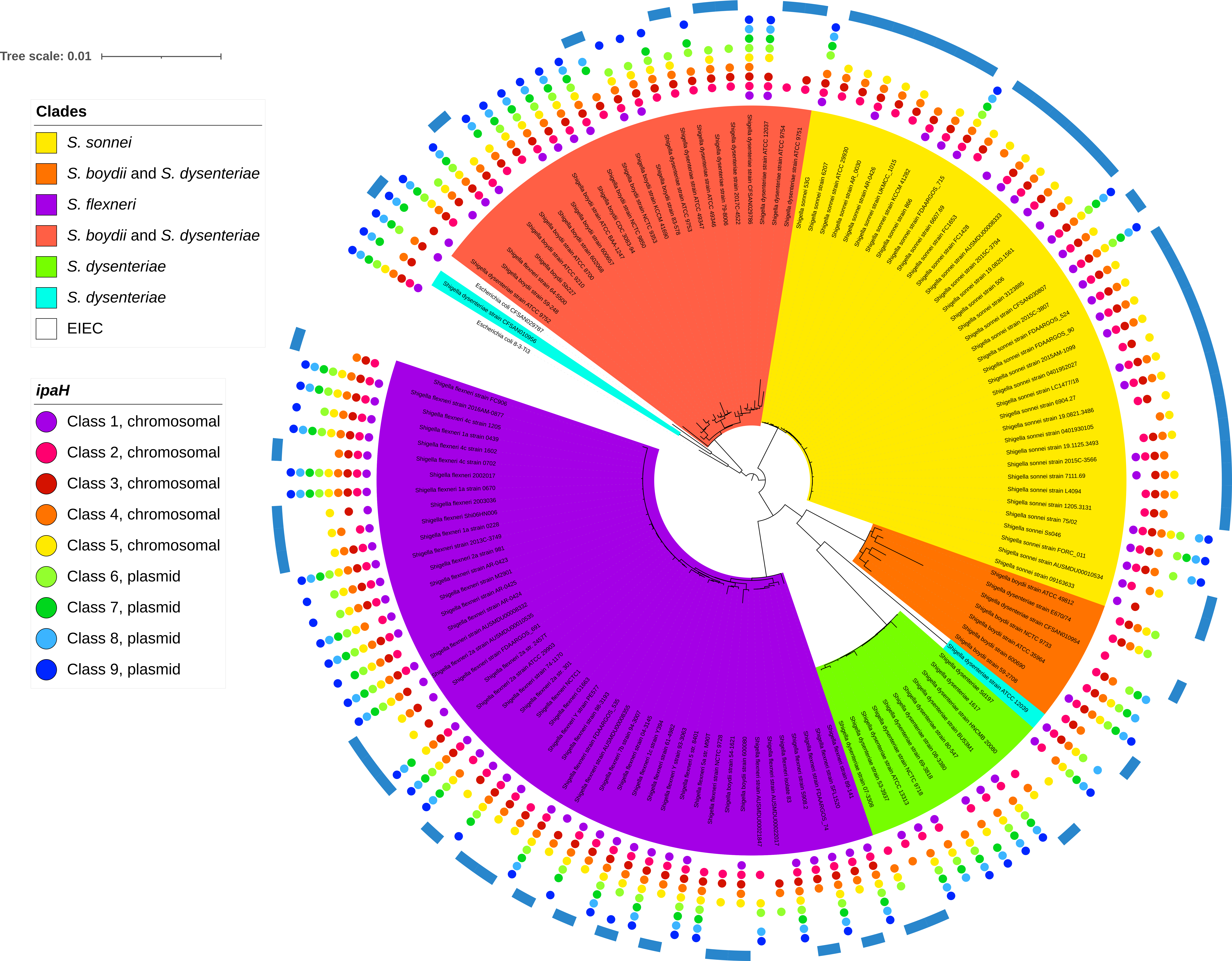

### Figure4.png

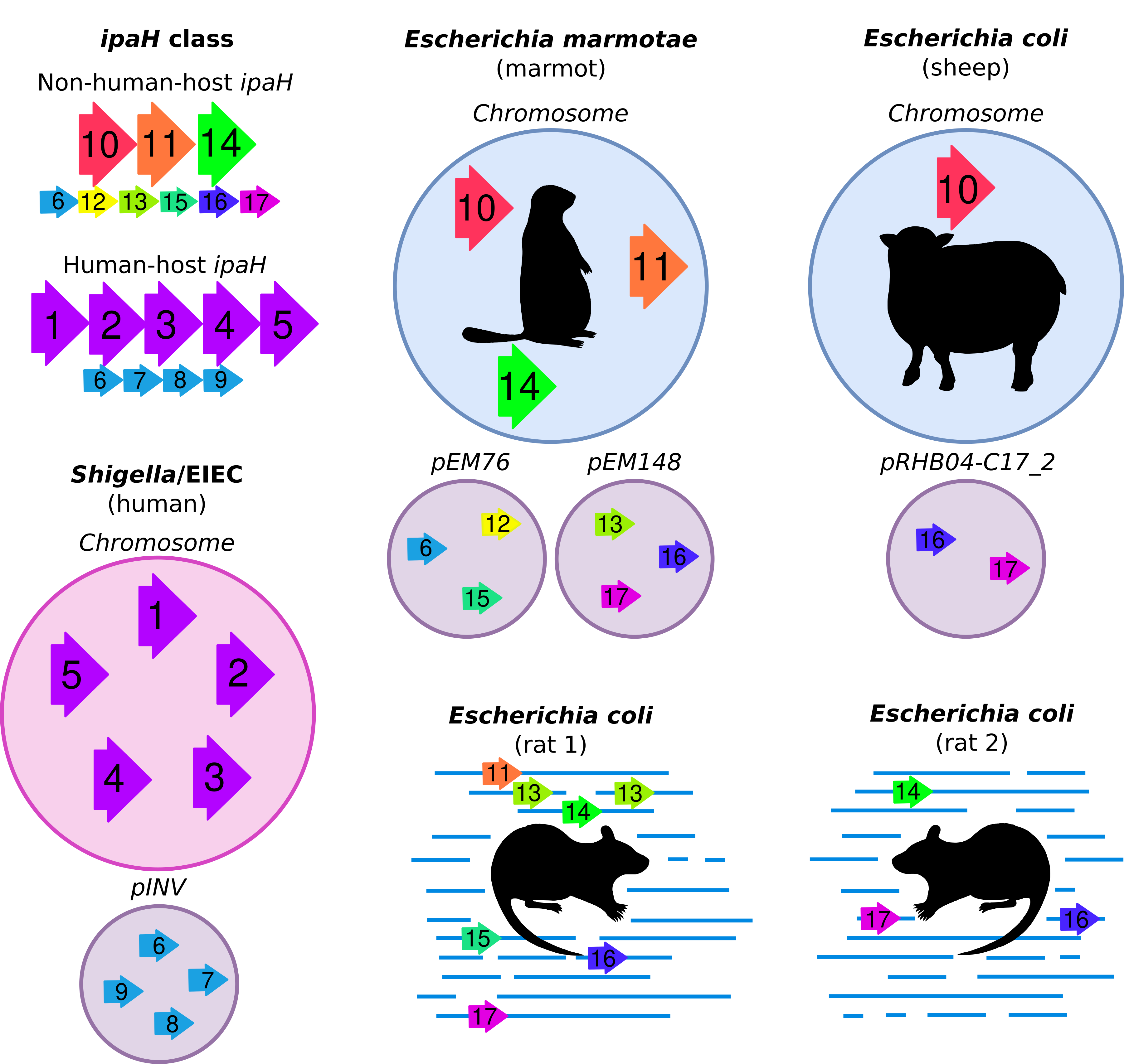

### Figure5.png

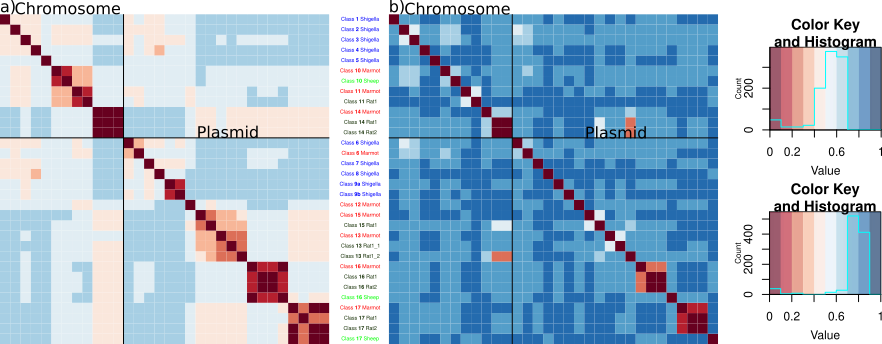

### Figure6.png

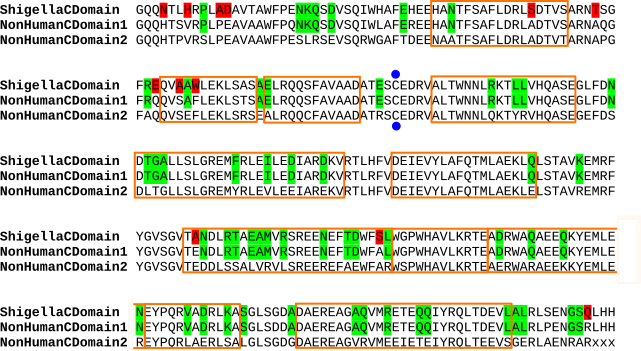

### SupplFigure1.png

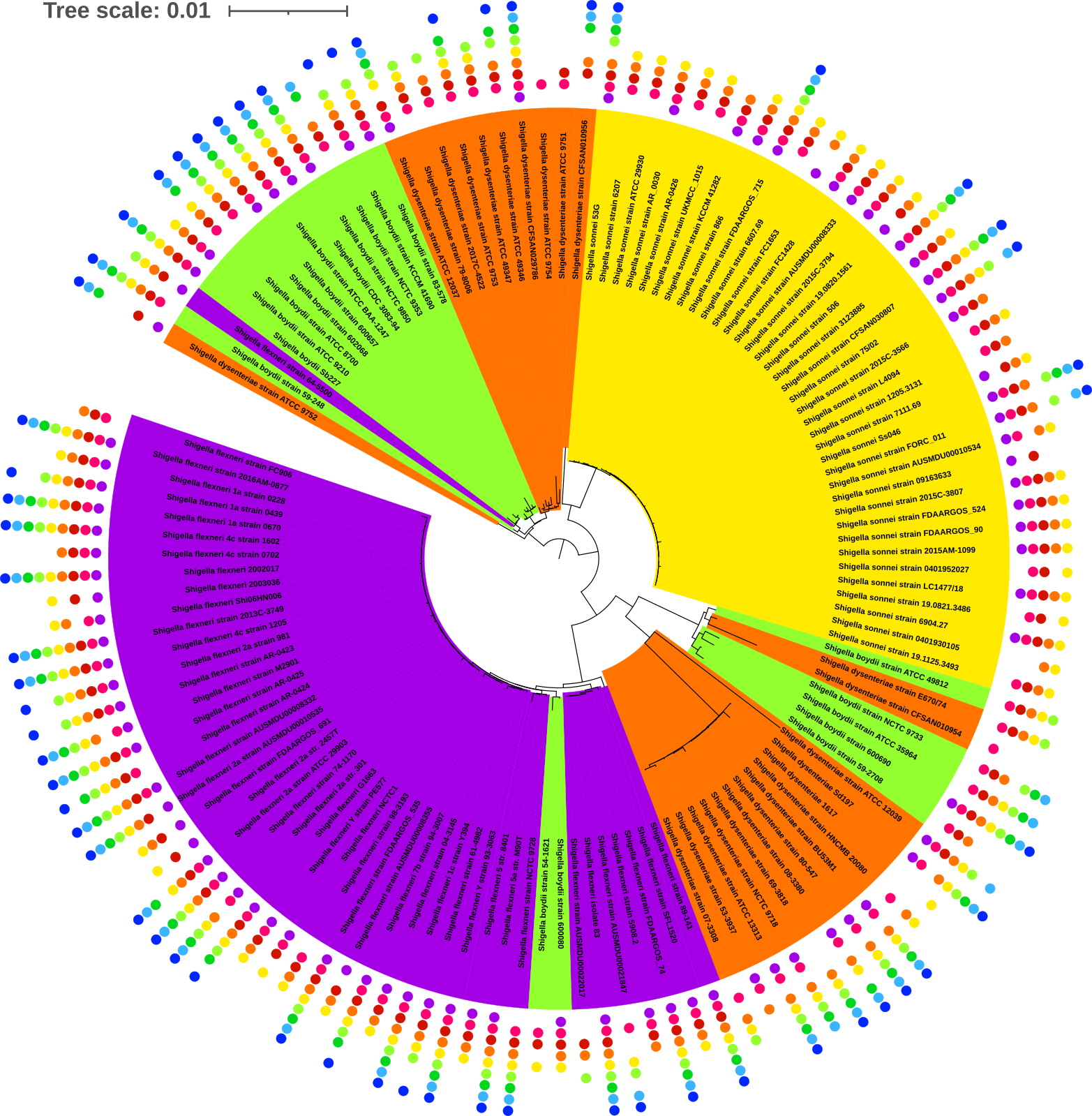

### SupplFigure2.png

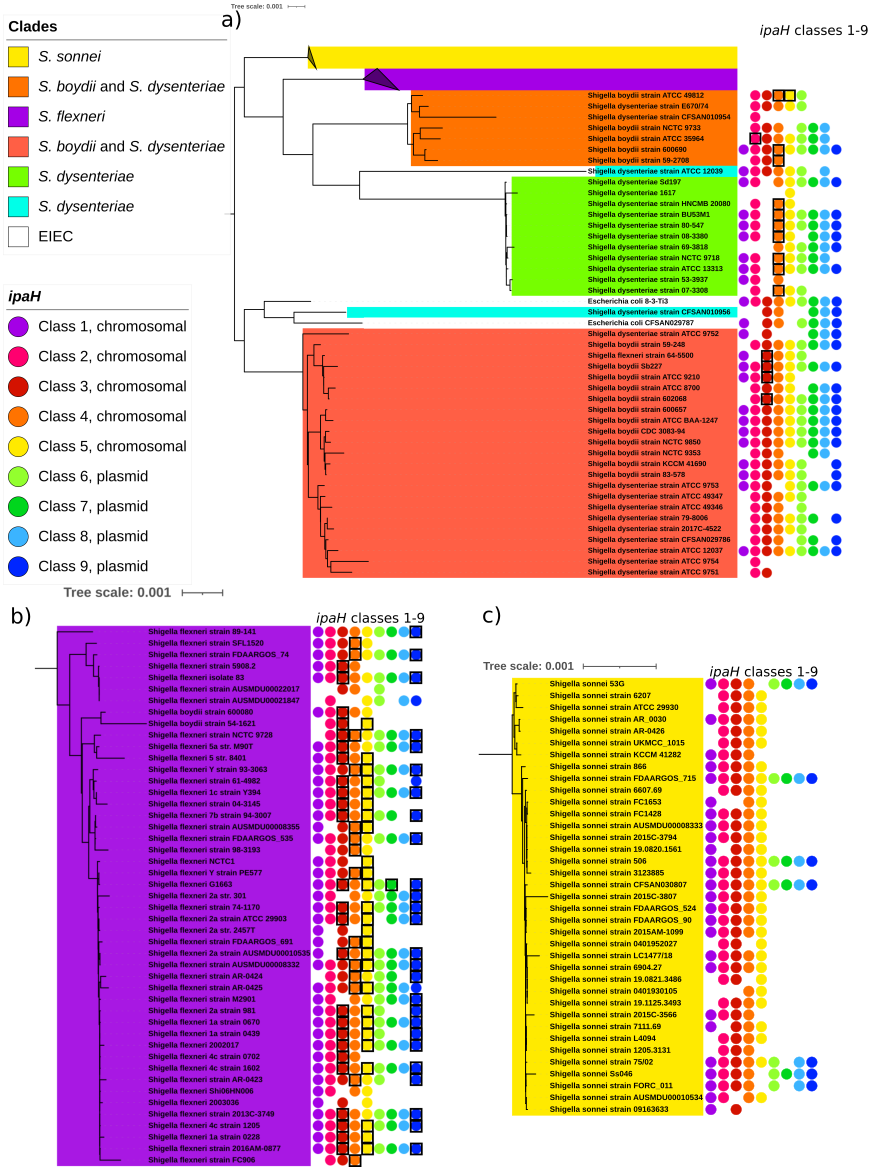

### SupplFigure3.png

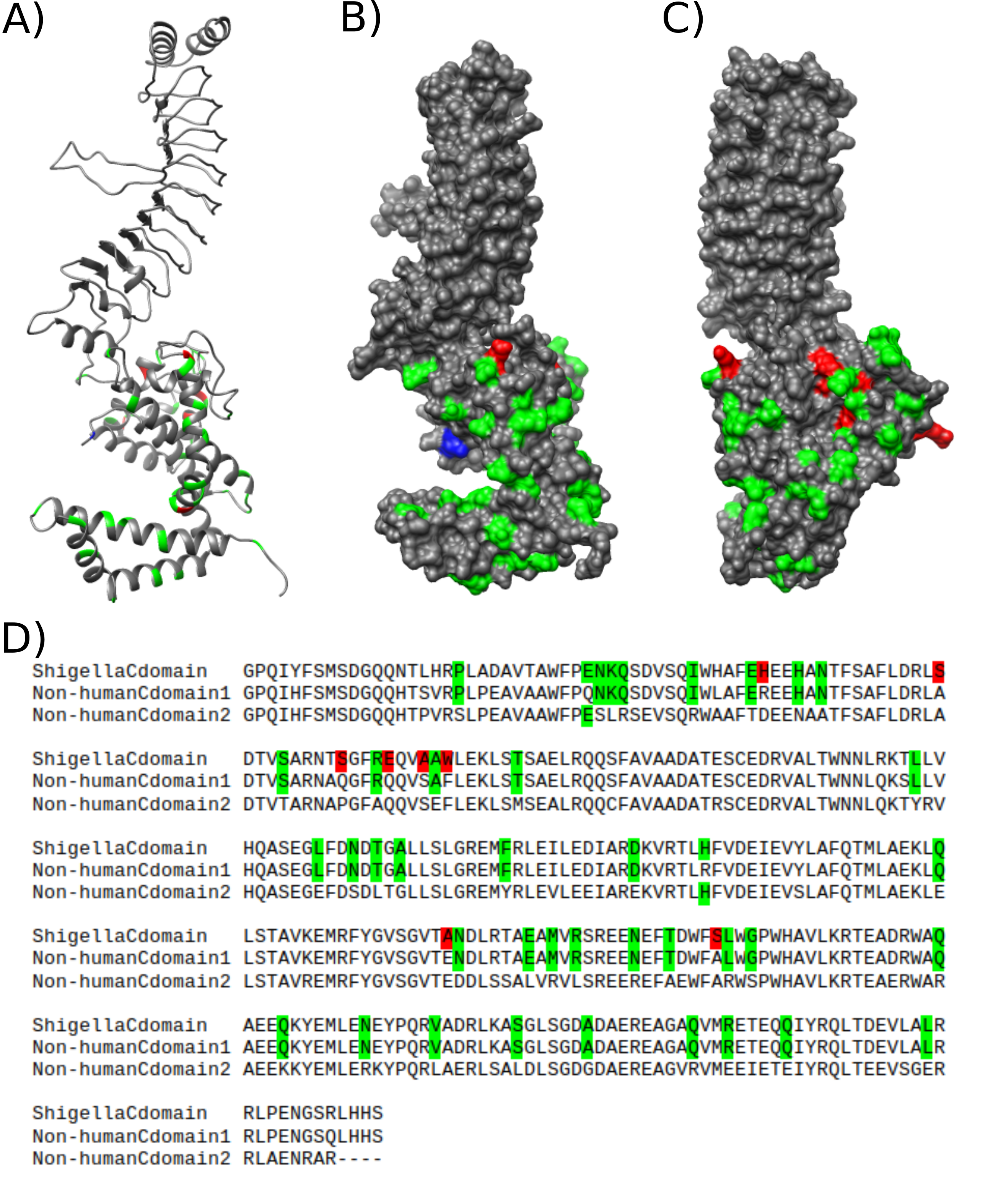
